# Dissecting Symptom-linked Dimensions of Resting-State Electroencephalographic Functional Connectivity in Autism with Contrastive Learning

**DOI:** 10.1101/2023.05.22.541841

**Authors:** Xiaoyu Tong, Hua Xie, Gregory A. Fonzo, Kanhao Zhao, Theodore D. Satterthwaite, Nancy Carlisle, Yu Zhang

## Abstract

Autism spectrum disorder (ASD) is a common neurodevelopmental disorder characterized by *social interaction deficits, communication difficulties*, and *restricted/repetitive behaviors or fixated interests*. Despite its high prevalence, development of effective therapy for ASD is hindered by its symptomatic and neurophysiological heterogeneities. To collectively dissect the ASD heterogeneity in neurophysiology and symptoms, we develop a new analytical framework combining contrastive learning and sparse canonical correlation analysis to identify resting-state EEG connectivity dimensions linked to ASD behavioral symptoms within 392 ASD samples. Two dimensions are successfully identified, showing significant correlations with social/communication deficits (r = 0.70) and restricted/repetitive behaviors (r = 0.45), respectively. We confirm the robustness of these dimensions through cross-validation and further demonstrate their generalizability using an independent dataset of 223 ASD samples. Our results reveal that the right inferior parietal lobe is the core region displaying EEG activity associated with restricted/repetitive behaviors, and functional connectivity between the left angular gyrus and the right middle temporal gyrus is a promising biomarker of social/communication deficits. Overall, these findings provide a promising avenue to parse ASD heterogeneity with high clinical translatability, paving the way for treatment development and precision medicine for ASD.

## Introduction

Autism spectrum disorder (ASD) is a common neurodevelopmental disorder that manifests in deficits in social interaction, communication, and restricted/repetitive behaviors or fixated interests^1^. ASD affects quality of life in both children and adults, yet treatment for ASD is limited. The heterogeneity in ASD, which includes variability in behavioral symptoms^2^, genetic profiles^3^, and neurobiology^4^, is a major obstacle to developing effective treatment. To address this heterogeneity, a better dissection of the psychopathological dimensions of ASD is needed to pave the way for the quantification of heterogeneity and potential effective treatment.

The core diagnostic criteria of ASD comprises two symptom domains: *social/communication deficits* (SCD), and *fixated interests or restrictive/repetitive behaviors* (RRB)^1^. Following these criteria, numerous studies have refined the ASD-relevant clinical behavior measures^5,6^ stemming from the gold standard tests for ASD^7,8^, aiming to construct a better representation of the symptomatic heterogeneity of ASD. However, the psychopathological essence of these pre-defined behavioral domains remains vague. For example, the RRB domains in the Autism Diagnostic Observation Schedule (ADOS)^7^ and Social Responsiveness Scale^9^ share the same name (both named as RRB), while only showing modest correlation with each other^10^. Encouragingly, recent work has suggested that machine learning techniques, such as sparse canonical correlation analysis (sCCA), can successfully establish data-driven symptom dimensions tailored to a neurophysiological basis^11^, providing a promising approach to circumvent limitations on symptom quantification imposed by clinical measures typically constrained by arbitrary diagnostic criteria. Furthermore, another recent study also established data-driven dimensions of ASD symptoms and identified potential subtypes based on brain functional connectivity associated with genotypes^12^, demonstrating the advantage of data-driven symptom dimensions over pre-defined behavior traits.

In addition to the symptomatic heterogeneity, neuropathological heterogeneity — the variability in underlying neurobiological alterations of ASD patients — is another important component of ASD heterogeneity hindering effective treatment development. Given the high variability observed across individual-level brain activity metrics, distilling disorder-specific information from overall variability would greatly facilitate the parsing of neuropathological heterogeneity in ASD. To this end, a machine learning technique termed contrastive learning, which extracts information that is unique in one dataset compared with the other, can be utilized to derive patient-specific neuroimaging features, thus eliminating the background information that is shared in both patients and typically developed (TD) subjects. Notably, a recent study employed a contrastive learning technique to identify principal neuroanatomical dimensions in ASD using structure-based neuroimaging data, demonstrating the potential for contrastive learning to reveal potential dimensions of neuropathology in ASD patients^13^. Together, these studies have provided evidence that neuroimaging data can be successfully used to establish data-driven symptom scores and links between symptoms and their corresponding neurophysiological bases.

However, all aforementioned studies have evaluated structural/functional magnetic resonance imaging (MRI) features. Despite its popularity and demonstrated utility in studies defining neural circuitry biomarkers^11,13,14^, the clinical utility of MRI is limited due to substantial requirements in terms of expertise, specialized equipment and prohibitive cost. Alternatively, electroencephalography (EEG) provides another neuroimaging data modality that is more cost-effective and easy-to-operate, which facilitates translation of research findings to improved clinical practice. In current study, using resting-state EEG (rsEEG) functional connectivity as neurophysiological features, we integrated contrastive learning with sCCA to parse the heterogeneity within ASD by identifying symptom-linked neurophysiological dimensions (Fig. 1).

**Fig. 1.**
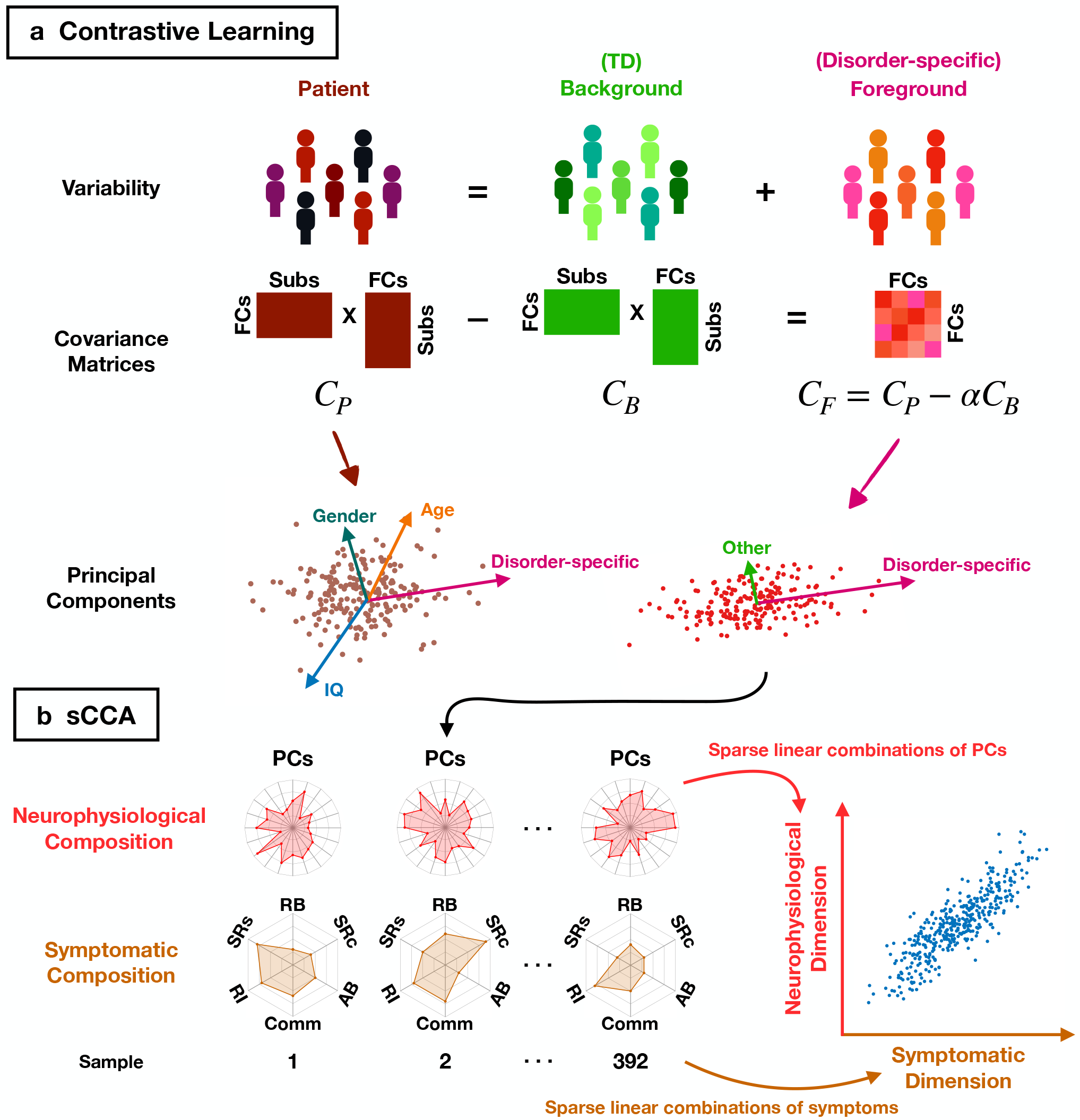
Flowchart of data-driven dissection of symptom-linked neurophysiological dimensions in ASD patients based on contrastive resting-state electroencephalographic functional connectivity (FC) features. **a** Contrastive Learning. Conceptually, the variability in ASD patients is composed of a background variability (shared by typically developing (TD) subjects) and a foreground variability (disorder-specific). The overall patient variability consists of not only disorder-related variance, but also the normative differences across subjects, such as age, gender, and intellectual capacity. To improve the signal-to-noise ratio of FC features, we extract contrastive features by subtracting the background variability from the overall patient variability. Specifically, we utilize covariance matrices to represent variability in subjects. Hence, the difference between (weighted) covariance matrices of ASD patients (C_P_) and TD subjects (C_B_) represents the foreground variability (C_F_), and is hypothetically related to mainly disorder-specific dimensions. Lastly, principal components (PC) are derived from the contrastive covariance matrix, and are used as the contrastive features for subsequent analyses. **b** Sparse Canonical Correlation Analysis (sCCA). sCCA is then conducted with the obtained contrastive features to correlate the neurophysiological profiles of ASD patients with their symptom traits, including repetitive behavior (RB), social reciprocity (SRc), adaptive behavior (AB), communication (Comm), retracted interests (RI), and social responsiveness (SRs). The linked dimensions between neurophysiology and symptoms are then identified, which are essentially the sparse linear combination of contrastive principal components and symptom subscales, respectively.

We identified two pairs of linked dimensions between neurophysiology and ASD symptoms, which respectively corresponded to SCD and RRB scales. We confirmed the robustness of these linked dimensions through cross-validation analysis and further demonstrated their generalizability using an independent dataset. Lastly but importantly, we further investigated the rsEEG functional connectivity signature associated with SCD and/or RRB symptoms. Together, our findings suggested two distinct brain mechanisms underlying SCD and RRB symptoms, providing a dissection of both symptomatic and neuropathological heterogeneities in ASD. This may more effectively motivate the development of effective ASD treatment and the achievement of precision medicine in clinical practice.

## Results

### Anchoring Latent Symptom Dimensions Using rsEEG Functional Connectivity

To extract disorder-specific features in ASD patients, we first performed contrastive principal component analysis (cPCA)^14^ using the rsEEG functional connectivity features (see Methods for details) on the Autism Biomarker Consortium for Clinical Trials (ABC-CT) dataset^15^ (N_TD_ = 239, N_ASD_ = 392). Specifically, the covariance of TD subjects quantified the subclinical (background) variability, which was then subtracted from the overall patient variability measured as the covariance of patient data. The resulting difference in covariance between overall patient variability and subclinical variability quantified the disorder-specific (foreground) variability, yielding contrastive functional connectivity features. To confirm the advantage of contrastive features, we conducted representational similarity analysis^16^ to assess the association between each type of connectivity features (standard, subclinical, contrastive) and clinical measures or demographic feature. As a result, we found that subclinical connectivity features were significantly more associated with age than contrastive features, demonstrating that the subclinical variability was indeed more closely associated with individual-level variance related to non-pathological brain characteristics (e.g. age). On the contrary, contrastive features were significantly more associated with symptom severity for most clinical measures compared with standard/subclinical features, suggesting the superiority of contrastive features in representing disorder-related variability in ASD patients (Supplementary Figs. 1, 2).

Afterward, sparse canonical correlation analysis (sCCA)^17^ was conducted to identify linked dimensions between the contrastive functional connectivity features and symptom profiles in ASD patients. To establish data-driven symptom scores for ASD, we incorporated a total of 24 subscales from Autism Diagnostic Observation Schedule (ADOS)^7^, Autism Impact Measure^6^, Social Responsiveness Scale^9^, and Vineland Adaptive Behavior Scale (VABS)^18^ to quantify the symptom profile of each patient (Supplementary Fig. 3). As a result, two pairs of generalizable linked dimensions were identified and cross-validated (ten-fold) between symptom profiles and contrastive functional connectivity features in THETA-band (correlation between symptom and functional connectivity dimensions: r = 0.70, the same correlation in cross-validation: r_CV_ = 0.16, permutation test for cross-validation: p_permutation_ = 0.005. Fig. 2a) and ALPHA-band (r = 0.45, r_CV_ = 0.28, p_permutation_ = 0.037. Fig. 2b), respectively.

**Fig. 2.**
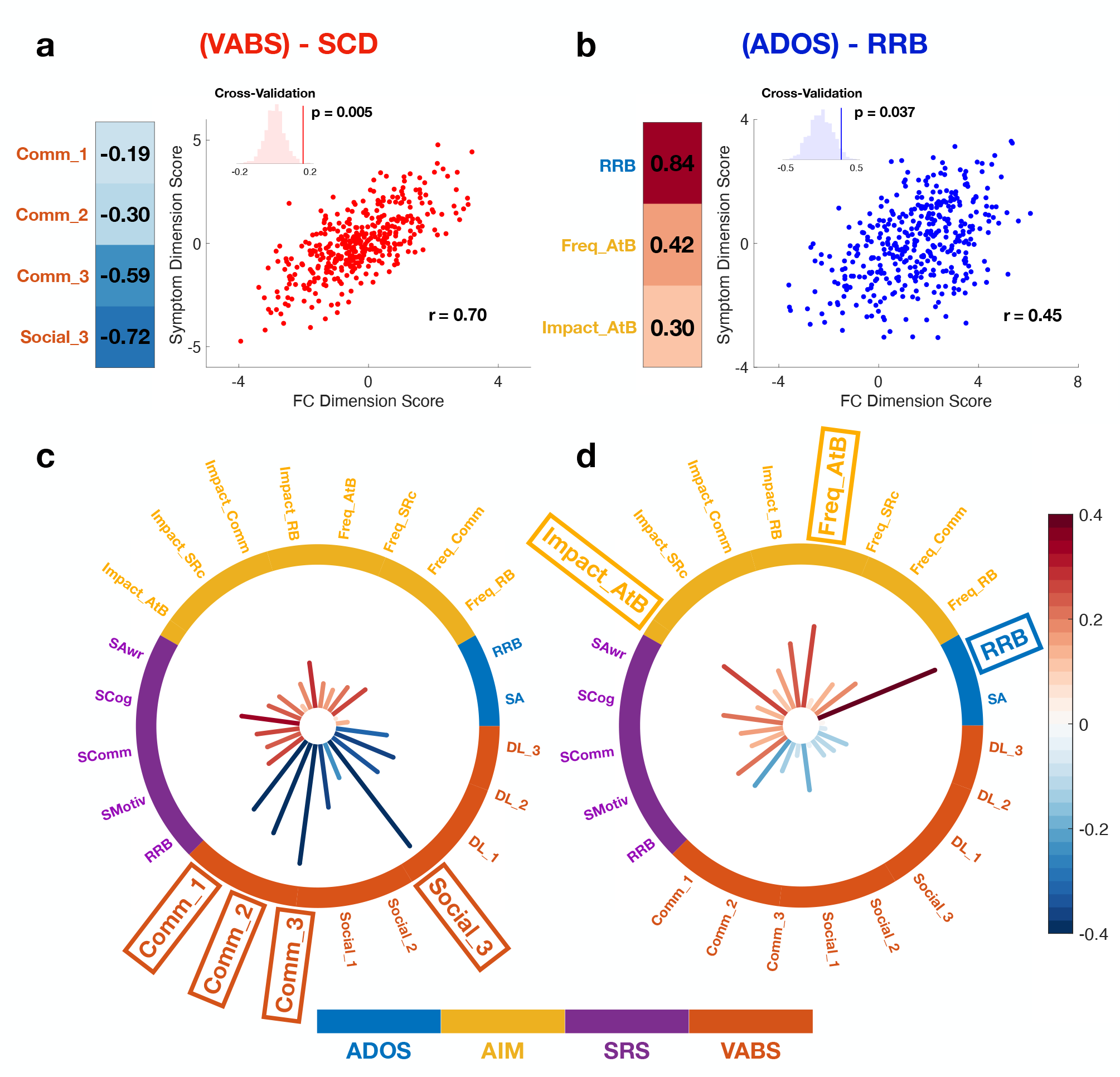
Linked dimensions between contrastive rsEEG features and symptom traits in ASD patients. **a** The social communication deficits (SCD) -linked dimension. The symptom dimension score is composed of the z-scores of deficits in communication (Comm_1: receptive; Comm_2: expressive; Comm_3: written) and socialization (Social_3: coping skills) functions as measured by VABS. The FC dimension score shows strong correlation with the symptom dimension score (r = 0.70) on ASD samples (n = 392). The robustness of the correlation between FC and symptom dimension scores is confirmed by cross-validation and permutation test (p_permutation_ = 0.005). The histogram in inset shows the null distribution of correlation coefficients between FC and symptom dimension scores over 1000 cross-validations, where the vertical line indicates the actual value without permutation (r_CV_ = 0.16). **b** The same figure as **a** with repetitive and restricted behavior (RRB) -linked dimension. The FC dimension score show strong correlation with the symptom dimension score (r_CV_ = 0.45) on ASD samples. The robustness of the correlation between FC and symptom dimension scores is confirmed by cross-validation and permutation test (p_permutation_ = 0.037). The histogram in inset shows the null distribution of correlation coefficient between FC and symptom dimension scores in cross-validation, where the vertical line indicates the actual value without permutation (r = 0.28). **c** Correlations between SCD-linked FC dimension scores and clinical measures. The FC dimension score shows strong negative correlation with the level of communication and socialization skills while only showing modest correlations with other clinical measures, demonstrating its specific association with SCD. **d** Correlations between RRB-linked FC dimension scores and clinical measures. The FC dimension score shows particularly strong positive correlation with the RRB severity, while showing relatively modest correlations with other clinical measures (including the impact and frequency (freq) of atypical behavior (AtB), which also contribute to the dimension score). This result demonstrates the specific association of this FC dimension score with SCD symptoms. **Abbreviations** for behavior scales: **ADOS**: Autism Diagnostic Observation Schedule; **AIM**: Autism Impact Measure; **SRS**: Social Responsiveness Scale; **VABS**: Vineland Adaptive Behavior Scale. For a complete list of clinical measures, see Supplementary Fig. 1.

The composition of symptom dimension score for THETA-band was attributed to communication and socialization subscales in VABS (Fig. 2a). Notably, VABS assesses the functionality of adaptive behaviors, thus being negatively correlated with the social and communication deficit (SCD). Therefore, greater dimension scores indicated more severe symptoms of SCD. Moreover, the symptom and functional connectivity dimension scores for THETA-band were strongly correlated with SCD as measured by behavior subscales in VABS (Fig. 2c, Supplementary Fig. 4a). These results suggest that the functional connectivity dimension identified in THETA-band is linked to SCD as quantified by VABS, thus is hereafter referred to as the SCD-linked dimension. Due to the intercorrelations between behavior scales in ASD patients (Supplementary Fig. 3), the symptom and functional connectivity dimensions also showed modest correlations with certain other behavior measures (Fig. 2c, Supplementary Fig. 4a).

In contrast, the composition of symptom dimension score for ALPHA-band was primarily driven by the RRB score in ADOS, with a small contribution from atypical behavior measured by Autism Impact Measure (Fig. 2b). Greater dimension scores were associated with more severe symptoms for RRB. Importantly, the symptom and functional connectivity dimension scores indeed showed strong correlations with ADOS-RRB (symptom: r = 0.74, p_FDR_ = 6.44 × 10^−69^. Supplementary Fig. 4b; functional connectivity: r = 0.40, p_FDR_ = 7.28 × 10^−5^. Fig. 2d). Similar to SCD-linked dimensions, the RRB-linked symptom and functional connectivity dimension scores also showed modest correlations with certain other behavior measures (Fig. 2d, Supplementary Fig. 4b) due to the significant correlations between incorporated behavior scales (Supplementary Fig. 3). Notably, these dimension scores did not show as strong correlations with the RRB score quantified by Social Responsiveness Scale, partly because of the modest correlation between ADOS-RRB and Social Responsiveness Scale-RRB (Supplementary Fig. 3). Overall, these results demonstrated that the functional connectivity dimension identified in ALPHA-band is linked to the RRB score as measured by ADOS, thus is hereafter referred to as the RRB-linked dimension.

We conducted further analyses to confirm that the identified SCD- and RRB-linked dimensions were not driven by demographic differences. Specifically, we found that both SCD- and RRB-linked symptom and functional connectivity dimensions were not significantly correlated with age (SCD: symptom: r = -0.084, p = 0.10; functional connectivity: r = -0.046, p = 0.36. RRB: symptom: r = -0.016, p = 0.76; functional connectivity: r = -0.023, p = 0.65. Supplementary Fig. 5), and there was significant group difference in dimension scores between genders (SCD: symptom: p_ranksum_ = 0.30; functional connectivity: p_ranksum_ = 0.38. RRB: symptom: p_ranksum_ = 0.16; functional connectivity: p_ranksum_ = 0.85. Supplementary Fig. 6). Moreover, we examined the correlation between SCD- and RRB-linked dimension scores. While no significant correlation was observed between SCD- and RRB-linked functional connectivity dimensions (r = 0.09, p = 0.07), a significant positive correlation was observed between the two identified symptom dimensions (r = 0.29, p = 4.13 × 10^−9^). These results suggested that the identified SCD- and RRB-linked functional connectivity dimensions (in THETA- and ALPHA-band, respectively) may correspond to distinct underlying mechanisms of ASD symptoms in different EEG frequency bands. Meanwhile, these functional connectivity dimensions may be associated with co-occuring symptoms that vary in severity and contribute to the heterogeneity across ASD patients.

### Functional Connectivity Signatures Associated with ASD Symptoms

Subsequently, we investigated the rsEEG functional connectivity signatures that constitute the symptom dimension of SCD and RRB. First, we examined the importance of network-level functional connectivity metrics (see **Methods** for details). For SCD, functional connectivity within the default mode network showed the highest importance (Fig. 3a). For RRB, functional connectivity within the default mode network and functional connectivity between the fronto-parietal control and visual networks showed the highest importance (Fig. 3b). Afterward, we further examined the importance of region-of-interest (ROI) level functional connectivity, as quantified by the feature weights for functional connectivity dimension loadings. Notably, functional connectivity between left angular gyrus and right middle temporal gyrus showed great importance for SCD (Fig. 4c), while the composition of the SCD-linked functional connectivity dimension was overall distributed across the fronto-parietal control network (Supplementary Fig. 7a). Meanwhile, we identified functional connectivity between left frontal eye field and right inferior parietal lobe as particularly important for the RRB-linked functional connectivity dimension (Fig. 4d). Among the top five most important ROI-level functional connectivity metrics for RRB, four of them involved right inferior parietal lobe (Fig. 4d, Supplementary Fig. 7b), indicating the strong association of alterations in right inferior parietal lobe’s neurophysiological activity wtih RRB symptoms.

**Fig. 3.**
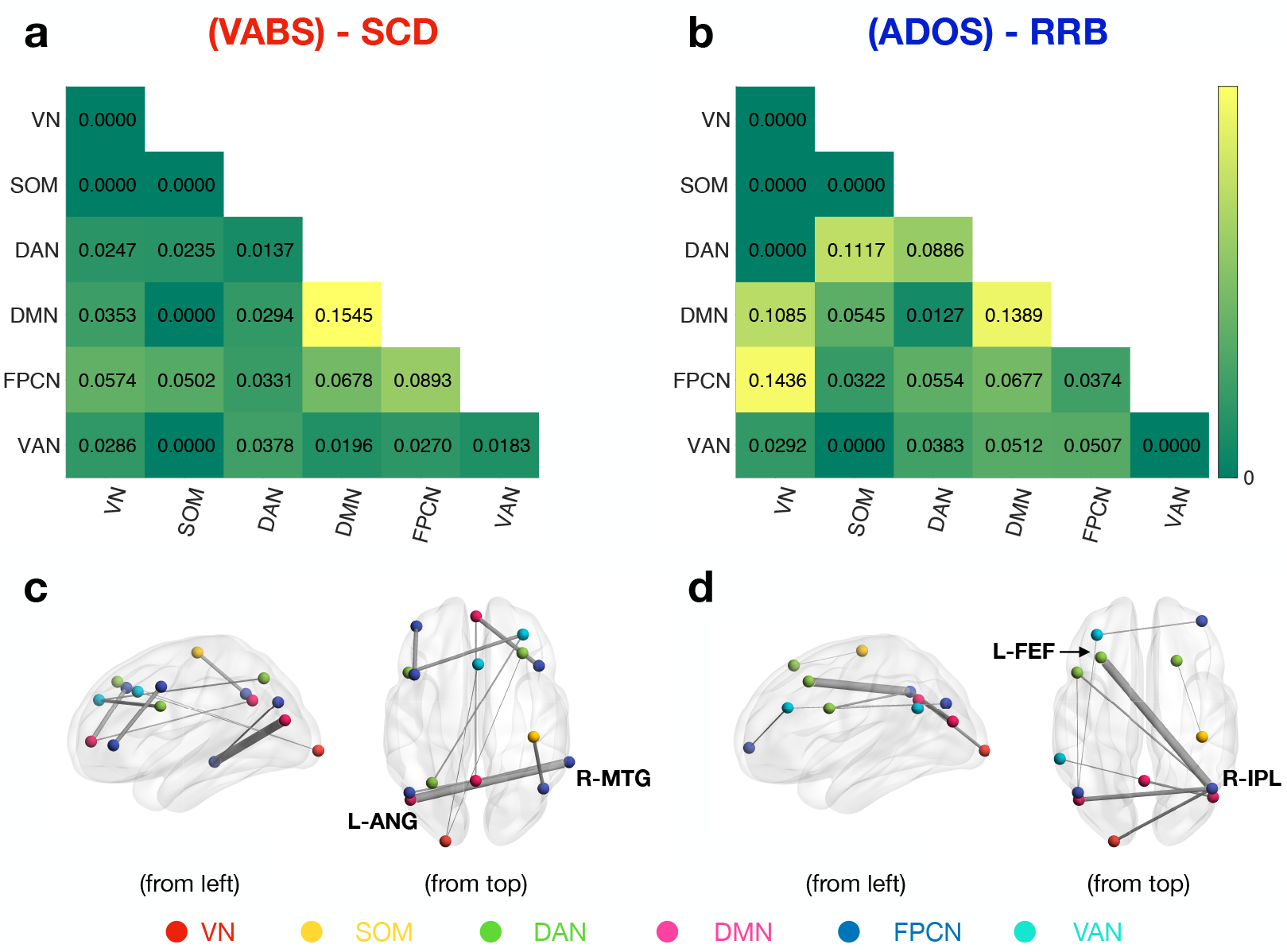
Signature alterations in rsEEG FC for ASD Symptoms. **a** Importance of network-level FC for SCD. The importance of network-level functional connectivity is calculated as the average absolute feature weights of all ROI-level functional connectivity within/between the brain network(s). The FC within DMN shows high importance for the severity of SCD. **b** Importance of network-level FC for RRB. Important FCs are more diffusive for RRB. The FCs between VN and FPCN and FCs within DMN show highest importance for RRB severity. Meanwhile, the FCs between DAN and SOM and FCs between DMN and VN also show a significant contribution to RRB severity. The importance of network-level FCs is derived by averaging the absolute feature weights of the top 10% strongest ROI-level FCs for better reliability and consistency. **c** Significant ROI-level FCs for SCD. The FC between left angular gyrus (**L-ANG**) and right middle temporal gyrus (**R-MTG**) shows the strongest association with SCD. **d** Significant ROI-level FCs for RRB. Right inferior parietal lobe (**R-IPL**) shows particularly strong association with the severity of RRB. Specifically, among the top five strongest ROI-level FC, four of them involve the R-IPL. The FC with strongest association with RRB severity is the one between R-IPL and left frontal eye field (**L-FEF**). The top 10 ROI-level FCs are shown for visualization purposes. **VN**: Visual Network. **SOM**: Somatomotor Network. **DAN**: Dorsal Attention Network. **DMN**: Default-Mode Network. **FPCN**: Fronto-Parietal Control Network. **VAN**: Ventral Attention Network.

**Fig. 4.**
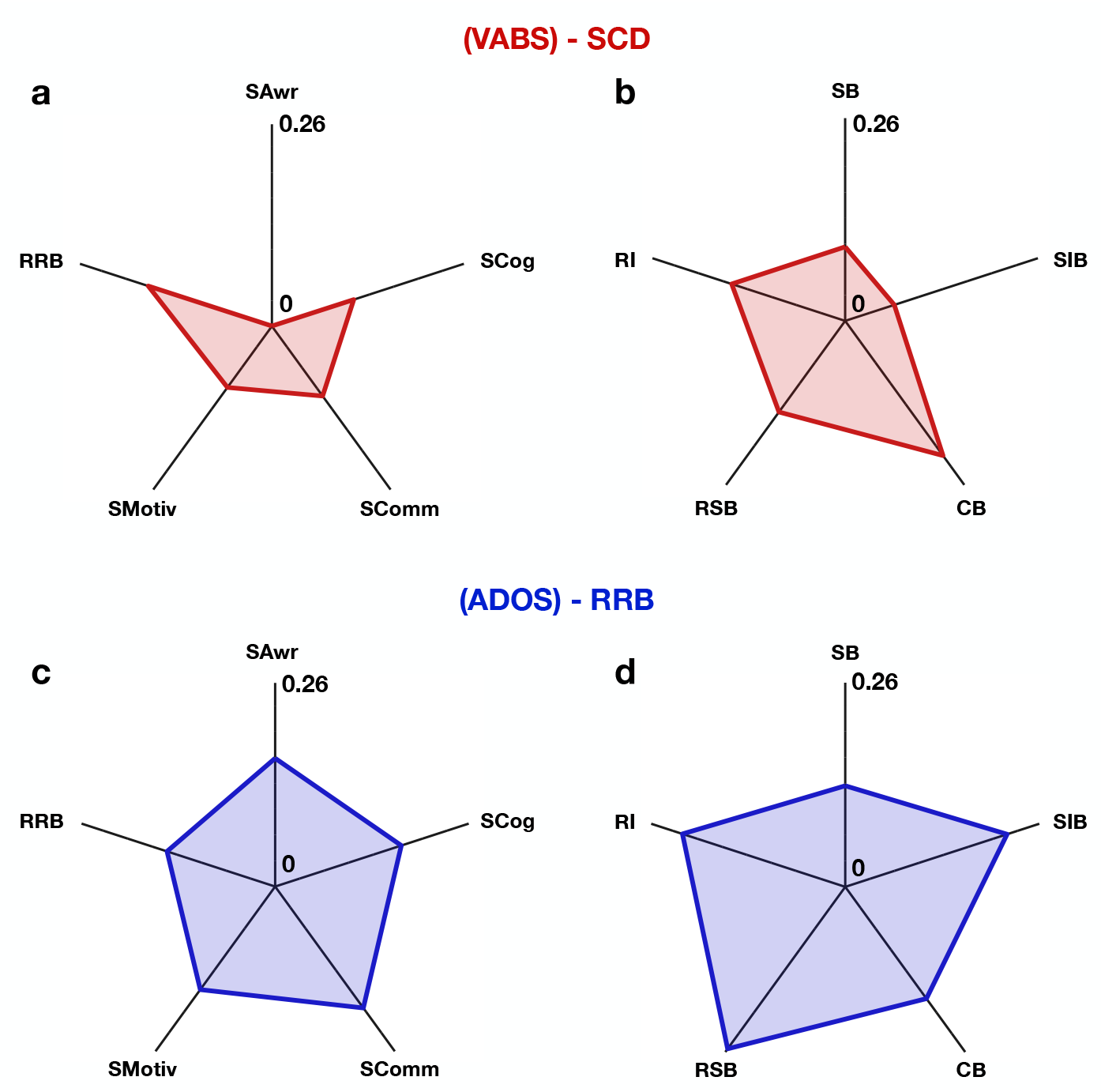
Association between symptom-linked FC dimensions and behavior traits in the independent HBN dataset. **a** Between SCD-linked FC dimension and Social Responsiveness Scale subscales. **b** Between SCD-linked FC dimension and Repetitive Behavior Scale subscales. **c** Between RRB-linked FC dimension and Social Responsiveness Scale subscales. **d** Between RRB-linked FC dimension and Repetitive Behavior Scale subscales. Symptom-linked dimensions show comparable (no significant difference) correlation strengths with Social Responsiveness Scale subscales between dataset 1 and the independent cohort, and also show significant positive correlations with Repetitive Behavior Scale subscales. Abbreviations: **SAwr**: Social Awareness; **SCog**: Social Cognition; **SComm**: Social Communication; **SMotiv**: Social Motivation; **SB**: Stereotypical Behavior; **SIB**: Self-Injurious Behavior; **CB**: Compulsive Behavior; **RSB**: Ritualistic/Sameness Behavior; **RI**: Restricted Interests.

### Generalization Analysis on an Independent Dataset

To confirm the generalizability of identified symptom-linked functional connectivity dimensions, we then conducted a validation analysis using an independent sample (N_ASD_ = 222) from the Healthy Brain Network (HBN) dataset^19^. We focused the generalization analysis on the functional connectivity dimensions, as the HBN dataset used a different set of behavior assessments. Specifically, we derived the transformation matrices (for SCD and RRB) from the ABC-CT dataset to convert standard rsEEG functional connectivity to the symptom-linked functional connectivity dimension scores. We then applied the same transformation matrices on HBN samples without further training and tuning to calculate the symptom-linked functional connectivity dimension scores on the independent dataset.

Subsequently, we compiled available ASD-related behavior scales in the HBN dataset with sufficient subjects (see **Methods** for details), including Social Responsiveness Scale and Repetitive Behavior Scale^20^. Afterward, we first examined the correlations between Social Responsiveness Scale subscales and symptom-linked functional connectivity dimension scores (Fig. 4a,c). As the Social Responsiveness Scale was the clinical measure shared by ABC-CT and HBN datasets, we then compared these correlation coefficients with the results for the ABC-CT dataset, which suggested that there was no significant difference in the correlation strengths when using the ABC-CT dataset or the HBN dataset (Fisher’s z test: SCD: social awareness: p_FDR_ = 0.29; social cognition: p_FDR_ = 0.07; social communication: p_FDR_ = 0.29; social motivation: p_FDR_ = 0.45; RRB: p_FDR_ = 0.94. RRB: social awareness: p_FDR_ = 0.94; social cognition: p_FDR_ = 0.31; social communication: p_FDR_ = 0.83; social motivation: p_FDR_ = 0.88; RRB: p_FDR_ = 0.12). These results suggest that the symptom-linked functional connectivity dimensions did not have substantially different correlative relationships with common behavior scales between cohorts.

Furthermore, to evaluate the generalizability of the association between identified functional connectivity dimensions and ASD symptoms, we examined the correlations between symptom-linked functional connectivity dimensions and subscales of Repetitive Behavior Scale on the HBN dataset (Fig. 4b,d), as subscales of Repetitive Behavior Scale have been shown to have significant correlations with both ADOS^21^ and VABS^22^. Encouragingly, we found that the SCD-linked functional connectivity dimension showed significant positive correlations with social cognition (r = 0.11, p = 0.048) and RRB (r = 0.15, p = 0.012) in Repetitive Behavior Scale. Again, it is worth reminding that there exists discrepancy between the RRB scores from different scales (Repetitive Behavior Scale vs. ADOS), which might affect the correlation of Repetitive Behavior Scale-RRB and SCD-linked dimension score. RRB-linked functional connectivity dimension also showed significant positive correlations with compulsive behavior (r = 0.15, p = 0.024) and restricted interests (r = 0.14, p = 0.031). Together, these results provide the generalizability of the association between symptom-linked functional connectivity dimensions and ASD symptom traits across unique samples and measures.

## Discussion

In this study, we identified two symptom-linked rsEEG functional connectivity dimensions in ASD patients by leveraging a combination of cutting-edge computational tools^11,14^. We showed that these two dimensions were mostly strongly-associated with SCD and RRB symptoms, two of the core symptom domains based on DSM-5^1,23^. Importantly, the symptom-linked rsEEG functional connectivity dimensions showed generalizable correlations with the corresponding ASD symptom dimension scores, as confirmed by cross-validation and a generalization analysis on an independent dataset, demonstrating initial evidence of reliability and generalizability. These findings provided a promising avenue for understanding symptomatic and neuropathological heterogeneity in ASD, which may guide the discovery of neuroimaging-informed treatment targets.

Despite the popularity and success of structural/functional MRI in identifying dimensions of brain structure/activity or psychopathology^11-14^, its clinical utility is limited due to substantial requirements in terms of expertise, specialized equipment, and high cost. We utilized rsEEG as an alternative data modality, which is more cost-effective and easy to operate. Additionally, thanks to its high temporal resolution, rsEEG functional connectivity can capture the typical/abnormal background oscillations (mainly alpha-band oscillation), which are highly indicative of ASD diagnosis^24^. Moreover, our method also enables the simultaneous identification of latent symptom dimensions and corresponding neurophysiological dimensions. Taken together, these innovations distinguish our current study from other existing work, demonstrating unique translational potential to improve clinical practice.

Notably, while we defined the neurophysiological dimensions as SCD- and RRB-linked, both SCD and RRB were composed of subscales^9,20,25,26^ and showed considerable inconsistency with themselves across assessments^21^. For example, both ADOS and Social Responsiveness Scale had RRB subscales, but they showed only modest correlations (r = 0.11, p = 0.023) in ASD patients on the ABC-CT dataset (N = 392). This discrepancy between subscales purporting to measure the same construct introduces additional heterogeneity to ASD symptom assessment. Therefore, we emphasize here that the RRB and SCD symptom dimension scores we constructed are specifically associated with the RRB score measured by ADOS and the SCD score composed by VABS. Moreover, as the identified ASD symptom dimension scores (SCD & RRB) were composed of loadings from multiple subscales, they represent data-driven dimensions unique from any existing measures. Therefore, we propose these new symptom dimensions be utilized in future research for quantifying ASD heterogeneity, as they exhibit optimal correlations with underlying neurophysiological features (See Fig. 2a,b for their compositions). If all involved psychiatric assessments are not available, we suggest approximating these symptom dimension scores using the loadings of corresponding measures from VABS and ADOS for best performance.

Remarkably, the symptom-linked rsEEG functional connectivity dimensions identified in this study are well-aligned with numerous previous studies of rsEEG abnormalities in ASD^24^. Specifically, we found that the decrease in functional connectivity between left angular gyrus and right middle temporal gyrus contributed most significantly to SCD-dimension symptom severity. While hypo-connectivity between left angular gyrus^27,28^ and right middle temporal gyrus^29-31^ has been widely reported to be associated with ASD diagnosis, our results offer more precise insights into their association with specific ASD symptom dimensions. For RRB, we found that the increase in functional connectivity between left frontal eye field and right inferior parietal lobe contributed substantially to RRB-domain symptom severity, echoing the findings of enhanced visual functioning^32^ and altered anatomical cortical networks in ASD^33^. We also identified that alpha-band right inferior parietal lobe activity was particularly important for RRB severity. Coinciding with our results, a previous pilot study^34^ applied repetitive transcranial magnetic stimulation (rTMS) over bilateral inferior parietal lobe to modulate alpha and low-beta band activities, and the authors found that the rTMS modulation differed according to the self-reported ASD symptom severity (as measured by Autism Spectrum Quotient^35^). These results support our findings that the right inferior parietal lobe may be a key region mediating RRB-domain severity and offer a potential treatment target that can be tested for alleviation of RRB symptoms.

Along with the findings from neuroimaging-based analyses, neurobiological studies have shown that disrupted development and connectivity of gamma-aminobutryic acid (GABA)ergic interneurons in the prefrontal and temporal cortices may be associated with ASD pathology^36,37^. Also, the alpha band is commonly associated with the active inhibition during rest^38,39^, a function achieved by GABAergic circuits^40^. Therefore, the alpha-band alterations in line with RRB severity could attribute to the malfunction of GABAergic circuits in the right inferior parietal lobe. Notably, Parkinson’s disease is another disorder that attributes to dysfunction of GABAergic circuit-based active inhibition, though the dysfunction is mostly proposed to be neurodegenerative instead of neurodevelopmental as in ASD. Deep brain stimulation is used as the first-line therapy for Parkinson’s disease^41,42^, but rTMS has also shown promising treatment outcomes^43^. In ASD, rTMS is preferred over deep brain stimulation for its non-invasive characteristics. Considering the aforementioned pilot study applying rTMS over bilateral inferior parietal lobe in ASD patients^34^, rTMS could be a promising potential therapy for ASD. Although other rTMS trials for ASD patients have shown less compelling results^44-46^, they applied rTMS to prefrontal regions, which might not be as effective a target as inferior parietal lobe, based on these and other findings. Also, previous studies did not consider heterogeneity in ASD, which may have reduced effect sizes. Taken together, we proposed that rTMS over inferior parietal lobe (specifically right inferior parietal lobe) might be a promising and effective therapy for ASD patients with high RRB traits.

In summary, we successfully parsed symptomatic and neurophysiological heterogeneity in ASD and provided initial evidence of generalizability with cross-validation and an independent dataset. We identified SCD and RRB as two principal ASD neurophysiology-linked symptom dimensions. Right inferior parietal lobe was identified as a key region associated with RRB symptom severity and might be targeted with rTMS as an intervention approach for ASD patients with high RRB traits. Functional connectivity between left angular gyrus and right middle temporal gyrus was the most highly-associated with SCD symptom severity, which might also be targeted by potential therapies targeting social impairment. Together, our findings provided a novel and exciting dissection of ASD heterogeneity, thus motivating discovery of novel, effective treatments and guiding precision medicine for ASD.

## Methods

### Dataset 1 — Autism Biomarker Consortium for Clinical Trials (ABC-CT) Dataset

#### Participants

The ABC-CT dataset^15^ comprised 280 children with ASD and 119 TD children whose ages ranged from 6 to 11. Each subject contained 1 to 3 EEG recordings and psychiatric assessments, resulting in 703 ASD samples and 364 TD samples. Due to low EEG data quality, 166 samples were excluded from our analysis (see **EEG Preprocessing** for details). Another 270 samples were also excluded because of incomplete psychiatric assessments, yielding 392 ASD samples and 239 TD samples that were included in our study. Diagnosis was made using the ADOS^7^. The Full-Scale Intelligence Quotient of subjects spanned from 60 to 150 as measured by Differential Ability Scales^47^. Exclusion criteria included known genetic syndrome, neurological condition putatively causally related to ASD, known metabolic disorder, and mitochondrial dysfunction. Medications were allowed to ensure the sample is representative to subjects with treatment for better generalizability of biomarker measurement, while a stable regimen (at least 8 weeks prior to enrollment) was required for inclusion.

#### Psychiatric Assessments

The ABC-CT dataset provided comprehensive measurements for ASD and TD subjects. The diagnostic characterization relied on ADOS^7^. While the Autism Diagnostic Interview-Revised^8^ scales were also proposed as diagnostic measurements^15^, they were excluded from our study due to incomplete data entries. Differential Ability Scales^47^ and VABS^18,26^ were included as clinician-administered assessments for intellectual and cognitive functionality. Additionally, this dataset also provided Autism Impact Measure^6^, Pervasive Developmental Disorder Behavior Inventory^48^, and Social Responsiveness Scale as caregiver questionnaires. However, the Pervasive Developmental Disorder Behavior Inventory was excluded from our study due to its limited data entries of only 50. The Clinical Global Impression Scale^49^ was also proposed to be employed as an outcome measure, but was not included due to lack of available data entries. Taken together, based on data availability and relevance to ASD, we included ADOS, VABS, Autism Impact Measure, and Social Responsiveness Scale as ASD-related symptom measures in our study.

Remarkably, the ABC-CT study employed a longitudinal design aligning with the structure and timeline of actual clinical trials^15^. Each subject was assessed at three time points (baseline, 6 weeks after baseline, and 24 weeks after baseline) for all aforementioned behavior measures, aiming for the evaluation of short-term test-retest reliability and developmental stability/change over a time period of typical clinical trials. Our results confirmed that the associations between symptom and functional connectivity dimension scores are general to all time points (Fisher’s z test for highest and lowest correlations. SCD: p = 0.28; RRB: p = 0.15. Supplementary Fig. 8). For information of distributions of demographics and psychiatric assessments for the ABC-CT dataset, see Supplementary Table 1.

#### EEG Acquisition

The ABC-CT studies employed four EEG paradigms, including resting-state (viewing abstract videos)^24^, human face event-related^50^, biological motion event-related^51^, and visual evoked^52^. In this study, we focused on the rsEEG features to establish neuroimaging biomarkers for ASD symptoms in naturalistic condition. Because the eye-closed session may not be reliably recorded for the subject age range (6-11), only eye-open sessions were collected. rsEEG data were recorded using a 128-channel EEG geodesic hydrocel system by Electrical Geodesics Inc. (EGI). The recordings were at a sampling rate of 1000 Hz. The recording reference was set to be Cz (head vertex) and the impedance of each electrode was checked before recording and was kept below 100 k*Ω*.

### Dataset 2 — Healthy Brain Network (HBN) Dataset

#### Participants

The HBN study recruited children and adolescents between 5–21 at four study sites in the New York City area^19,53^. The recruitment criterion was having adequate verbal communication ability with the help of parents or guardians. Exclusion criteria included severe neurological disorders, severe impairment in cognition (IQ < 66), acute encephalopathy, known neurodegenerative disorder, or other abnormalities that may prevent full participation in the protocol. Overall, HBN incorporated a variety of psychiatric disorders, and we focused on ASD in this study. At the time we accessed the dataset, it included 185 TD subjects and 222 ASD patients (regardless of comorbidity) with available behavior measures and neuroimaging data (one for each subject).

#### Psychiatric Assessments

The HBN dataset provided comprehensive assessments for diagnostic psychiatry, intelligence, language, and other cognitive functions in a transdiagnostic population^19^. In this study, we focused on the ASD-related assessments and intellectual capacity, as intellectual capacity has been shown to be associated to heterogeneity in psychiatric disorders^54^ and the ABC-CT dataset also contained comparable assessments (DAS). Although ADOS was proposed as the diagnostic assessment in the HBN dataset^19^, it was unavailable at the time we acquired the data. We included the Social Responsiveness Scale and Repetitive Behavior Scale^20^ for the evaluation of ASD symptoms on the HBN dataset. Notably, the Gendered Autism Behavioural Scale^55^ and VABS were also related to the generalization analysis, but there were not enough available data entries (no data was available for Gendered Autism Behavioural Scale; only 24 ASD patients with both available VABS assessments and EEG data). Additionally, subjects aged 6-17 were also assessed with Wechsler Intelligence Scale for Children^56^ for their intellectual capacity. Supplementary Table 2 provides information on the distributions of demographics and psychiatric assessments for the HBN dataset.

#### EEG Acquisition

On the contrary to the psychiatric assessments, EEG data was collected at a single study site^19^. For the resting state paradigm, which was designed to measure intrinsic brain activity during rest^57^, participants were asked to view a fixation cross on the center of the computer screen and to open or close their eyes at various points as instructed. High-density EEG data were recorded in a sound-shielded room using a 128-channel EEG geodesic hydrocel system by Electrical Geodesics Inc. (EGI). The recordings were at a sampling rate of 500 Hz with a bandpass of 0.1 to 100 Hz, and the recording reference was set to be Cz (head vertex). The impedance of each electrode was checked before recording and was kept below 40 k*Ω*.

### EEG Preprocessing

The recorded resting-state EEG data were cleaned offline with a fully automated artifact rejection pipeline built upon the EEGLab toolbox^58^. The entire procedure included the following steps: (1) The resampling of EEG to 250 Hz. (2) The removal of 60 Hz a.c. line noise artifact^59^. (3) The removal of non-physiological low frequency in the EEG signals using a 0.01 Hz high-pass filter. (4) The rejection of bad epochs by thresholding the magnitude of each epoch. (5) The rejection of bad channels by thresholding the spatial correlations among channels. (6) The exclusion of subjects with more than 20% bad channels. (7) The estimate of EEG signals from bad channels from the adjacent channels via the spherical spline interpolation^60^. (8) An independent component analysis to remove remaining artifacts, including scalp muscle artifact, ocular artifact, and ECG artifact^61^. (9) Re-referencing EEG signals to the common average. (10) The filtering of EEG signals to four canonical frequency ranges: theta (4–7 Hz), alpha (8–12 Hz), beta (13–30 Hz), and low gamma (31–50 Hz).

### EEG Source-Space Functional Connectivity Calculation

With the Brainstorm toolbox^62^, we first implemented source localization using the minimum-norm estimation approach^63^ to convert the channel-space EEG into the source-space signals of 3,003 vertices. A three-layer (scalp, skull, and cortical surface) boundary element head model was computed with the OpenMEEG plugin^64^ based on the FreeSurfer average brain template^65^. A total of 3,003 dipoles with unconstrained orientations were generated on the cortical surface. The lead-field matrix relating the dipole activities to the EEG was obtained by projecting the standard electrode positions onto the scalp. For each subject, an imaging kernel that maps from the channel space EEG to the source space current density was then estimated by the minimum norm estimation approach with depth weighting and regularization. Principal component analysis was then employed to reduce the three-dimensional estimated source signal at each vertex to the one-dimensional time series of the principal component.

To capture the brain functional architecture, we extracted connectomic features by calculating power envelope connectivity (PEC)^66^ since it has demonstrated strength in mitigating spurious correlations resulting from volume conduction^67^. Hilbert transform was first applied to convert source estimates into analytical time series. The analytical time series of each pair of brain signals were then orthogonalized to remove the zero-phase-lag correlation^67^. Afterward, the power envelopes were measured by calculating the square of the orthogonalized analytical signals. A logarithmic transform was subsequently conducted to enhance normality. PEC was then calculated as the Pearson’s correlation coefficient between the log-transformed power envelopes of each pair of brain regions. Finally, a Fisher’s r-to-z transformation was performed to enhance normality^67,68^. The regional pairwise PEC features were further extracted for the 31 regions of interest (ROIs) in the MNI (Montreal Neurological Institute) template space^69,70^. For each pair of regions, connectivity was calculated by averaging PEC values over all possible vertex pairs.

### Extraction of Contrastive Functional Connectivity Features for ASD Patients

To improve the specificity of functional connectivity features to ASD symptoms, we extracted contrastive functional connectivity features using the contrastive principal component analysis (cPCA) technique^14^. Specifically, 239 TD samples in the ABC-CT dataset were employed as the background data to identify subclinical variability in functional connectivity features shared across TD subjects. Meanwhile, 392 ASD samples in the ABC-CT dataset were utilized to identify variability in functional connectivity features shared across ASD patients. The cPCA was jointly conducted on both groups to quantify representative dimensions for subclinical and patient variabilities. As the variability in ASD patients can be conceptualized as the superposition of subclinical and ASD-specific variabilities, contrastive functional connectivity features were quantified as the difference between patient and subclinical principal components of functional connectivity features. Lastly but importantly, some subclinical variability might share the same directions as ASD-indicative dimensions — they were subclinical only because TD subjects had smaller magnitude in those dimensions. Therefore, a contrastive parameter (*α*, between 0 and 1) was included to control the quantity of subtracted subclinical variability. The contrastive parameter was optimized using cross-validation for each EEG condition with increment of 0.1. Supplementary Figs. 1 and 2 show the comparison across standard, subclinical, and contrastive principal components for THETA- and ALPHA-band EEG functional connectivity, respectively.

### Sparse Canonical Correlation Analysis (sCCA)

CCA is a multivariate procedure that seeks maximal correlations between linear combinations of variables in a pair of data matrices^71^. Sparsity constraint was applied by L1-norm regularization to reduce the model complexity and alleviate the overfitting issue^17^. sCCA was employed to identify generalizable linked dimensions between contrastive rsEEG functional connectivity and ASD-related symptom scores. Specifically, contrastive principal components of rsEEG functional connectivity yielded by cPCA was first scaled to have a unit variance. This step was to ensure the sparsity constraint is fair for all input features. No de-mean step was implemented after the cPCA step to make sure the overall procedure (including cPCA and sCCA steps) could be formularized as a linear transformation, to facilitate generalization analysis on independent dataset. Symptom measures were also normalized to have a zero mean and unit variance before the sCCA procedure.

### Grid Search for Hyperparameters with Cross-Validation

There were three hyperparameters required for the entire analysis, including one contrastive parameter used in cPCA, two sparsity parameters used in sCCA for contrastive functional connectivity features and symptom scores, respectively. These hyperparameters were optimized simultaneously within 5 rounds of 10-fold cross-validation. For the cross-validation, the correlation between canonical dimensions of contrastive functional connectivity features and symptom scores was used as the metrics for the evaluation. The samples from same subject were either all in the training set or all in the test set.

### Generalizability Analysis on Independent Dataset

For the generalization analysis of symptom-linked dimension scores, we first multiplied the cPCA loading, scaling for unit variance, and sCCA loading together to derive the equivalent linear transformation to convert standard rsEEG functional connectivity features to symptom-linked functional connectivity dimension scores for both RRB and SCD. Afterward, we directly applied the same linear transformation on the rsEEG functional connectivity features in the HBN dataset, where the rsEEG functional connectivity features were also normalized for each subject as in the ABC-CT dataset. We then conducted correlation analysis on the derived functional connectivity dimensions with the symptom scores on the HBN dataset to test the aforementioned hypotheses. Both datasets were normalized for each subjects before the procedure.

### Importance of Network-Level Functional Connectivity

In principle, the importance of network-level functional connectivity is calculated as the average absolute feature weights of all ROI-level functional connectivity within/between the brain network(s). To improve the reliability, we only included the top 10% significant ROI-level functional connectivity to derive the importance of network-level functional connectivity, as the contribution of ROI-level functional connectivity with small loadings for the dimension score could be non-significant and unstable.

## Acknowledgements

This work was supported by NIH grant nos. R21MH130956, R01MH129694, and Lehigh University FIG (FIGAWD35), CORE, and Accelerator grants. Portions of this research were conducted on Lehigh University’s Research Computing infrastructure partially supported by NSF Award 2019035. GAF was supported by NIH grant nos. K23MH114023 and R01MH125886 and grants from the Brain and Behavior Research Foundation and One Mind – Baszucki Brain Research Fund. TDS was supported by NIH grant nos. R37MH125829, R01EB022573, R01MH112847, and R01MH120482.

## Notes

### Competing Interest Statement

The authors have declared no competing interest.

