## Supplementary Material for "Dissecting Symptom-linked Dimensions of Resting-State Electroencephalographic Functional Connectivity in Autism with Contrastive Learning"

### Supplementary Materials

**Supplementary Figure 1** Comparison across standard, subclinical, and contrastive principal components for THETA-band EEG FC
**Supplementary Figure 2.** Comparison across standard, subclinical, and contrastive principal components for ALPHA-band EEG FC
**Supplementary Figure 3.** Correlations across ASD-related clinical measures in ASD patients
**Supplementary Figure 4.** Correlations between symptom dimension scores and clinical measures.
**Supplementary Figure 5.** Correlations between age and FC/symptom dimension scores
**Supplementary Figure 6.** Distribution differences between gender and FC/symptom dimension scores
**Supplementary Figure 7.** Composition of ROI-level FC for symptom-linked FC dimensions.
**Supplementary Figure 8.** Associations between symptom and FC dimension scores are general to longitudinal assessments

**Supplementary Table 1** Distribution of Demographics and Psychiatric Assessments of ASD Patients in ABC-CT Dataset

**Supplementary Table 2** Distribution of Demographics and Psychiatric Assessments of ASD Patients in HBN Dataset

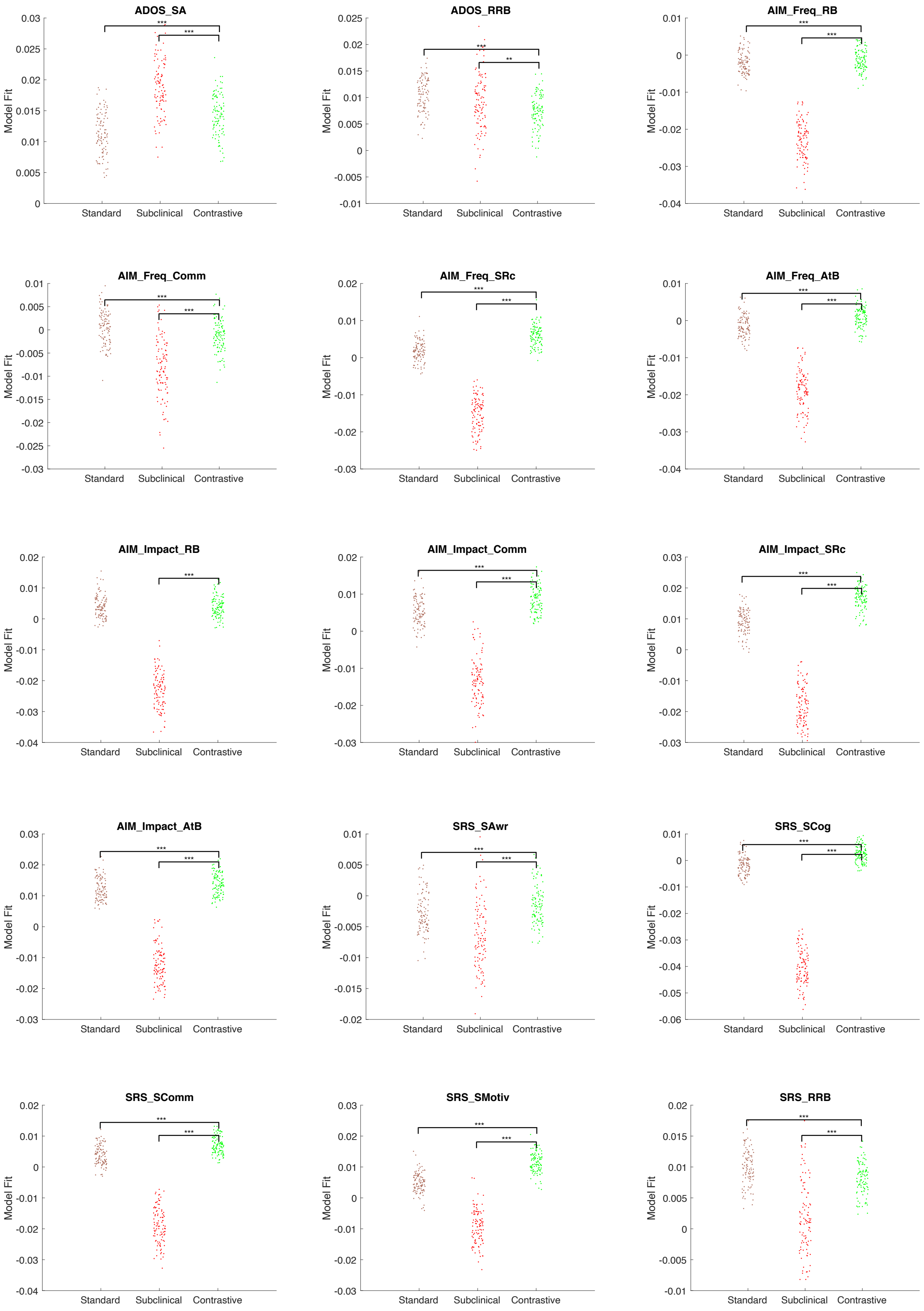

ADOS_SA
ADOS_RRB
AIM_Freq_RB
AIM_Freq_Comm
AIM_Freq_SRc
AIM_Freq_AtB
AIM_Impact_RB
AIM_Impact_Comm
AIM_Impact_SRc
AIM_Impact_AtB
SRS_SAwr
SRS_SCog
SRS_SComm
SRS_SMotiv
SRS_RRB
Model Fit
Standard
Subclinical
Contrastive
***
**

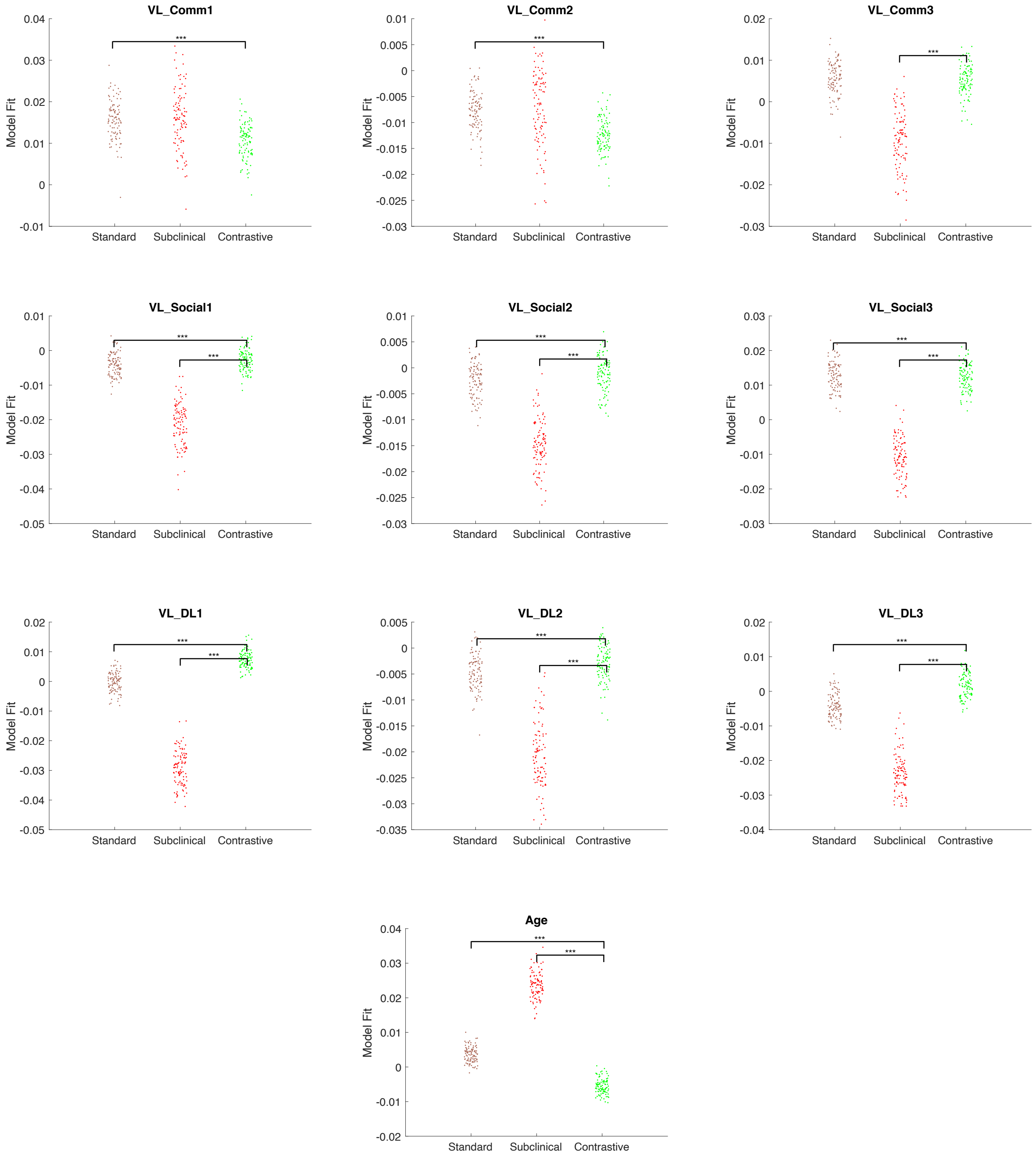


**Supplementary Figure 1.** Comparison across standard, subclinical, and contrastive principal components for THETA-band EEG FC. Representational similarity analysis (with Spearman's rho) is conducted to assess the association between FC features and each of the clinical measures. The significance level is determined by Wilcoxon's signed-rank test. p-values are corrected using FDR method. *** $p < 0.001$. ** $p < 0.01$. * $p < 0.05$.

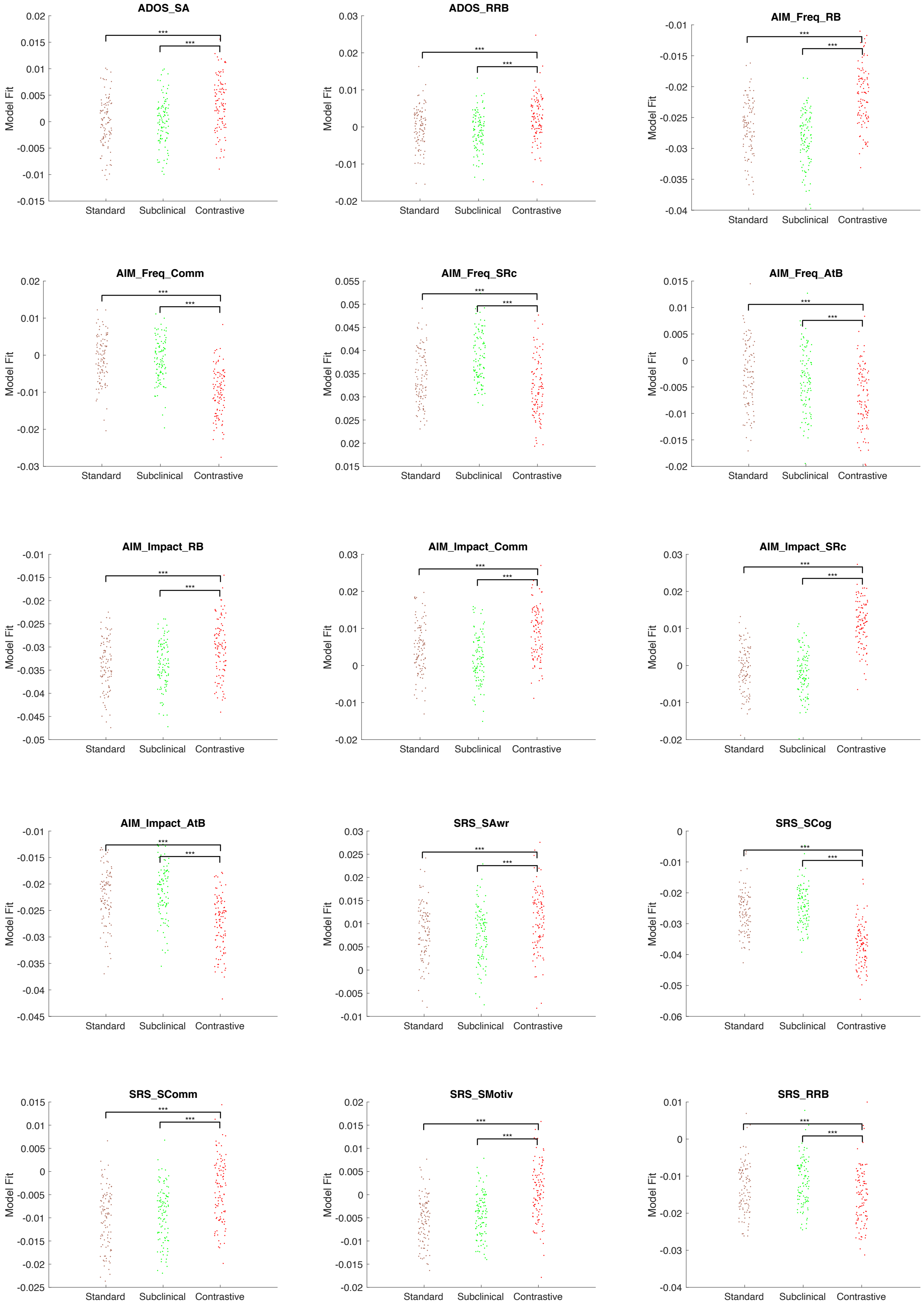
ADOS_SA
0.02
0.015
0.01
0.005
0
-0.005
-0.01
-0.015
Model Fit
***
***
Standard
Subclinical
Contrastive
ADOS_RRB
0.03
0.02
0.01
0
-0.01
-0.02
Model Fit
***
***
Standard
Subclinical
Contrastive
AIM_Freq_RB
-0.01
-0.015
-0.02
-0.025
-0.03
-0.035
-0.04
Model Fit
***
***
Standard
Subclinical
Contrastive
AIM_Freq_Comm
0.02
0.01
0
-0.01
-0.02
-0.03
Model Fit
***
***
Standard
Subclinical
Contrastive
AIM_Freq_SRc
0.055
0.05
0.045
0.04
0.035
0.03
0.025
0.02
0.015
Model Fit
***
***
Standard
Subclinical
Contrastive
AIM_Freq_AtB
0.015
0.01
0.005
0
-0.005
-0.01
-0.015
-0.02
Model Fit
***
***
Standard
Subclinical
Contrastive
AIM_Impact_RB
-0.01
-0.015
-0.02
-0.025
-0.03
-0.035
-0.04
-0.045
-0.05
Model Fit
***
***
Standard
Subclinical
Contrastive
AIM_Impact_Comm
0.03
0.02
0.01
0
-0.01
-0.02
Model Fit
***
***
Standard
Subclinical
Contrastive
AIM_Impact_SRc
0.03
0.02
0.01
0
-0.01
-0.02
Model Fit
***
***
Standard
Subclinical
Contrastive
AIM_Impact_AtB
-0.01
-0.015
-0.02
-0.025
-0.03
-0.035
-0.04
-0.045
Model Fit
***
***
Standard
Subclinical
Contrastive
SRS_SAwr
0.03
0.025
0.02
0.015
0.01
0.005
0
-0.005
-0.01
Model Fit
***
***
Standard
Subclinical
Contrastive
SRS_SCog
0
-0.01
-0.02
-0.03
-0.04
-0.05
-0.06
Model Fit
***
***
Standard
Subclinical
Contrastive
SRS_SComm
0.015
0.01
0.005
0
-0.005
-0.01
-0.015
-0.02
-0.025
Model Fit
***
***
Standard
Subclinical
Contrastive
SRS_SMotiv
0.02
0.015
0.01
0.005
0
-0.005
-0.01
-0.015
-0.02
Model Fit
***
***
Standard
Subclinical
Contrastive
SRS_RRB
0.01
0
-0.01
-0.02
-0.03
-0.04
Model Fit
***
***
Standard
Subclinical
Contrastive

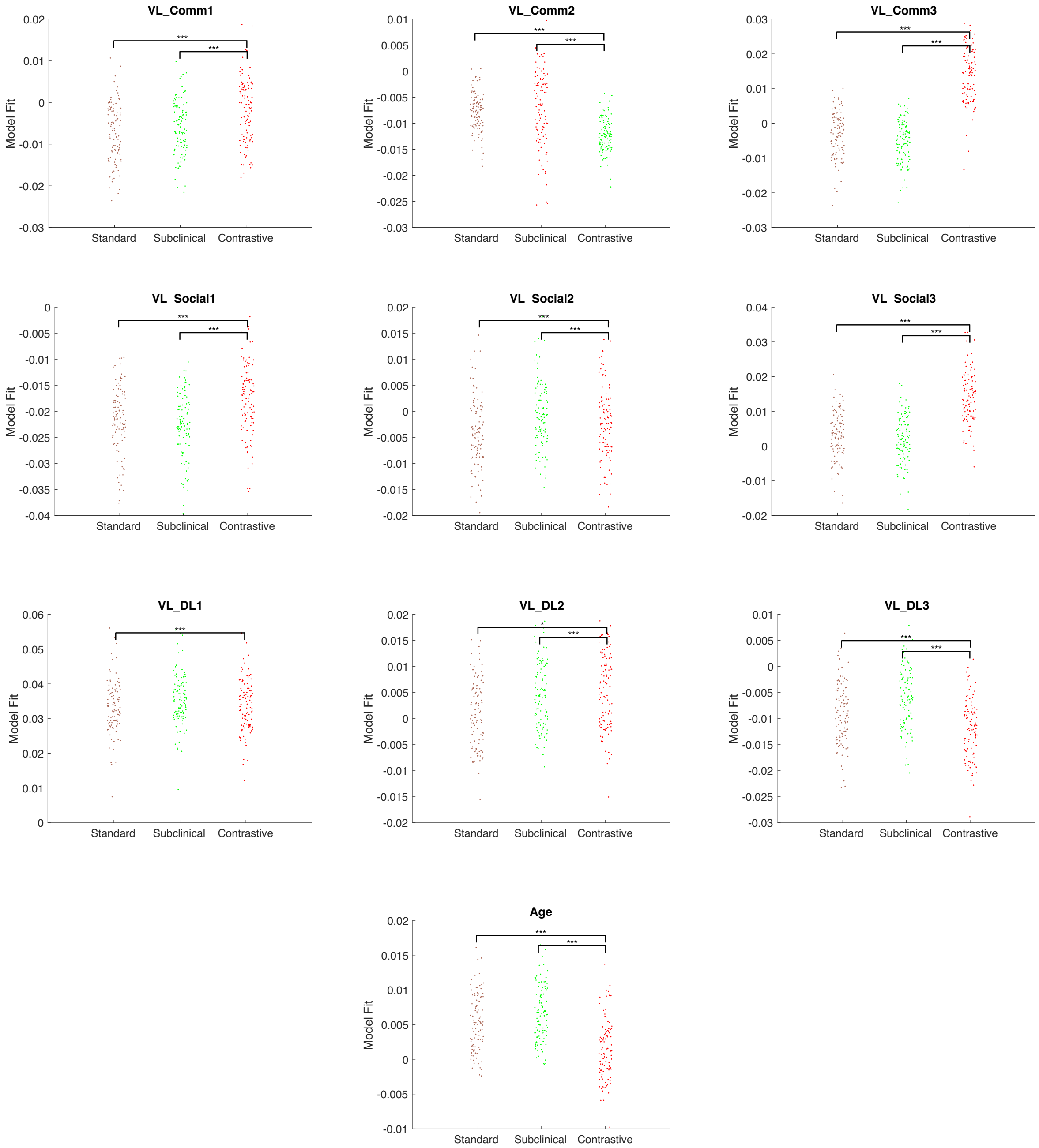


**Supplementary Figure 2.** Comparison across standard, subclinical, and contrastive principal components for ALPHA-band EEG FC. Representational similarity analysis (with Spearman's rho) is conducted to assess the association between FC features and each of the clinical measures. The significance level is determined by Wilcoxon's signed-rank test. p-values are corrected using FDR method. *** $p < 0.001$. ** $p < 0.01$. * $p < 0.05$.

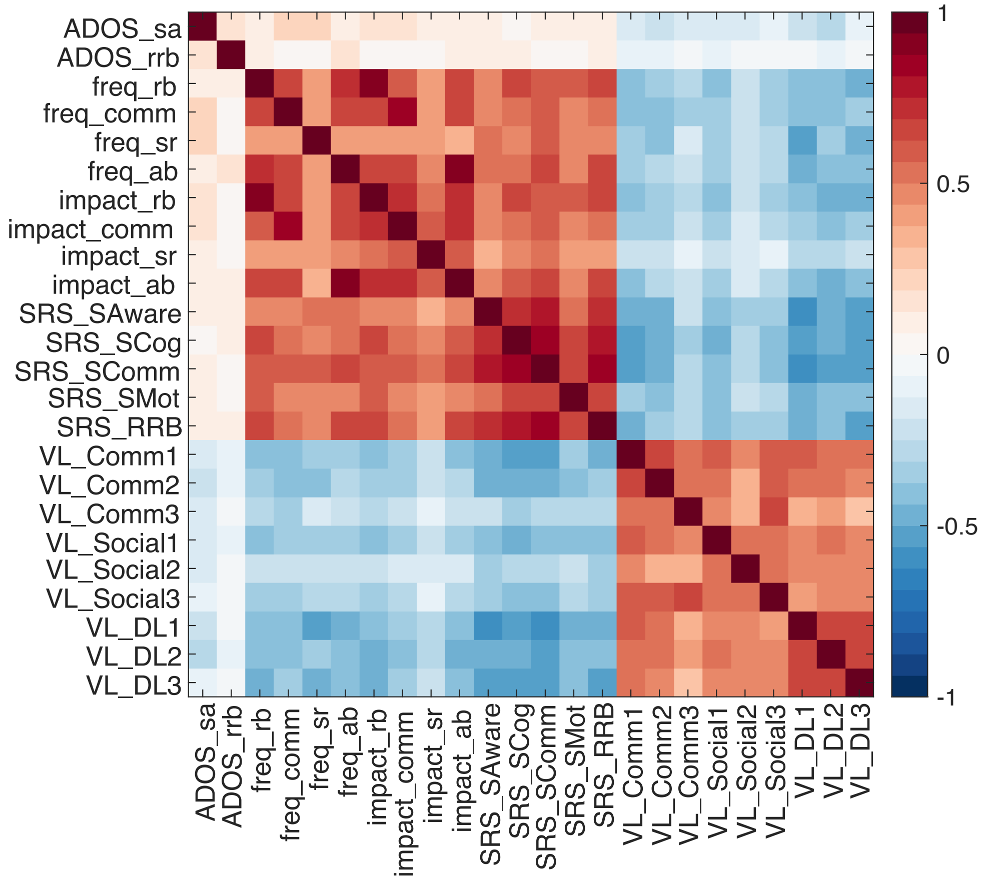


**Supplementary Figure 3.** Correlations across ASD-related clinical measures in ASD patients. Abbreviations: **ADOS**: Social Affect, Restrictive and Repetitive Behaviors; **AIM**: Frequency & Impact: Repetitive Behaviors, Communication Language, Social Reciprocity, Atypical Behavior; **Social Responsiveness Scale**: Social Awareness, Social Cognition, Social Communication, Social Motivation, Restricted Interests and Repetitive Behavior; **VABS**: Communication (receptive, expressive, written), Socialization (interpersonal relationships, play and leisure, coping skills), Daily Living (person, domestic, community)

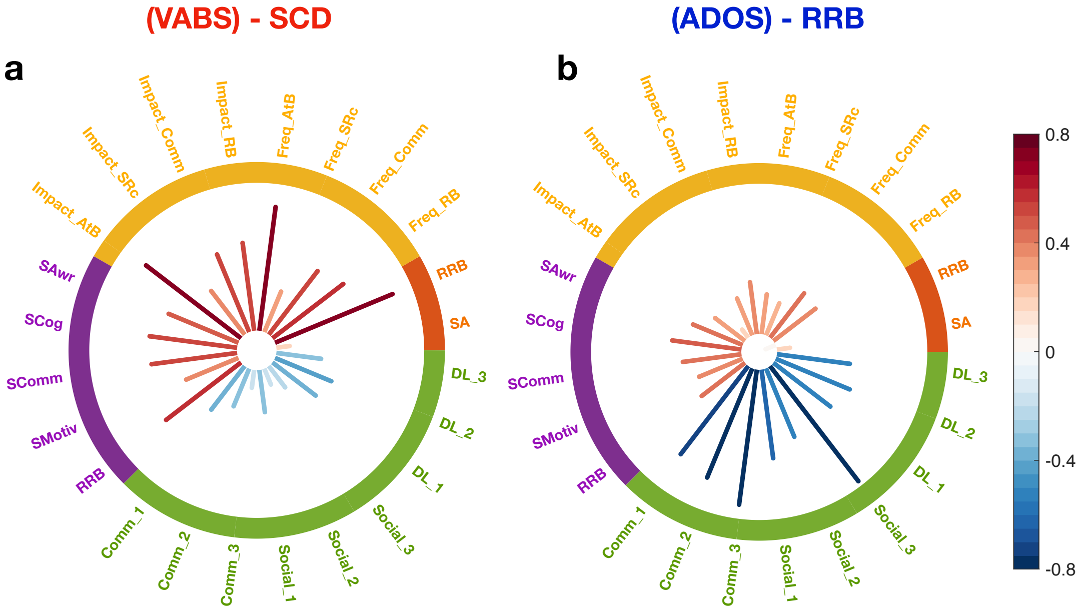


**Supplementary Figure 4.** Correlations between symptom dimension scores and clinical measures. **a** SCD-dimension. The symptom dimension score shows strong negative associations with social and communication skills, namely strong positive associations with social and communication deficits. **b** RRB-dimension. The symptom dimension score shows strong associations with clinical measures for RRB and atypical behavior.

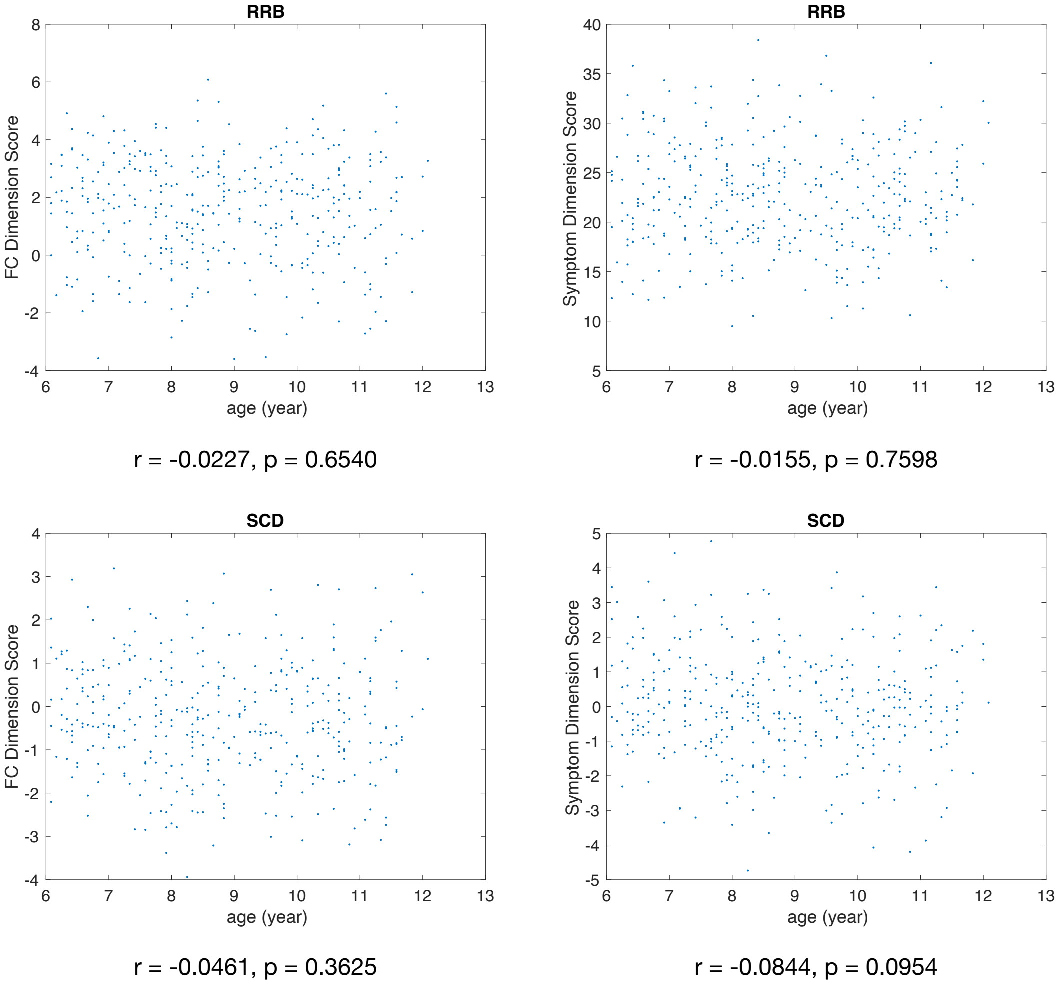


**Supplementary Figure 5.** Correlations between age and FC/symptom dimension scores. FC/symptom dimension scores for both RRB and SCD show independence of age. p-values are uncorrected.

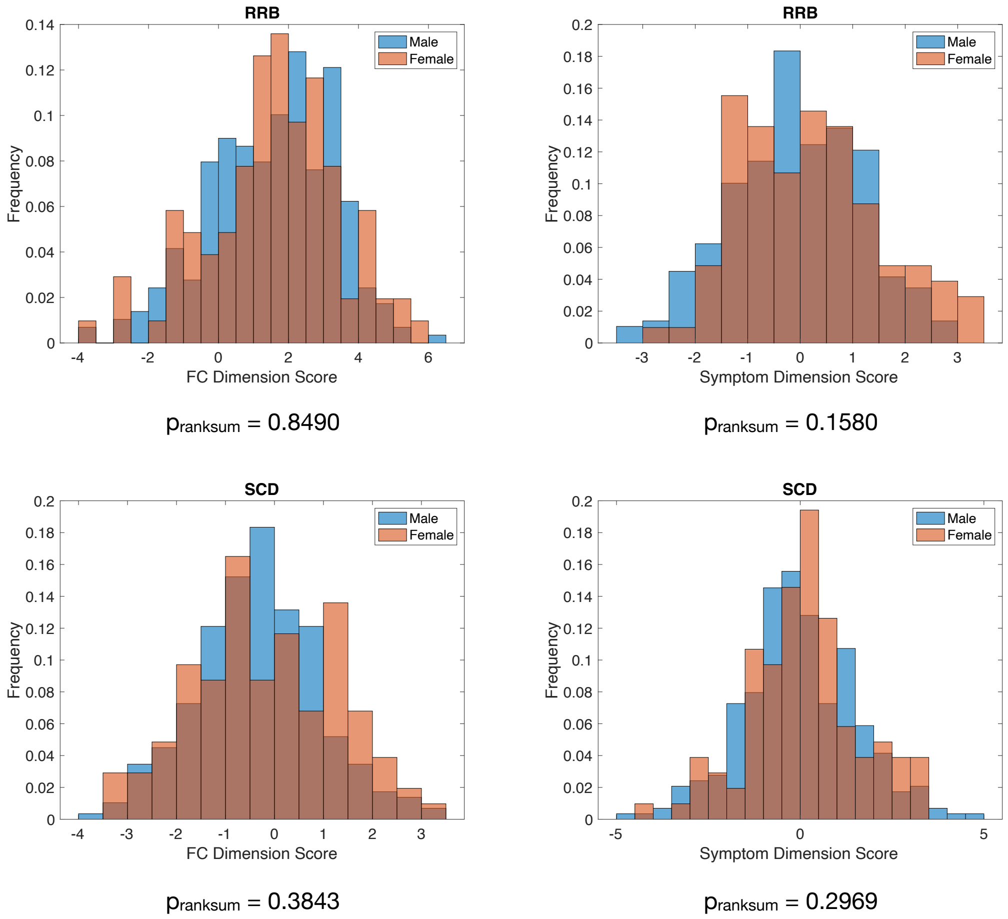


**Supplementary Figure 6.** Distribution differences between gender and FC/symptom dimension scores. FC/symptom dimension scores for both RRB and SCD show independence of gender. p-values are calculated based on Wilcoxon's ranksum test on dimension scores between male and female groups. p-values are uncorrected.

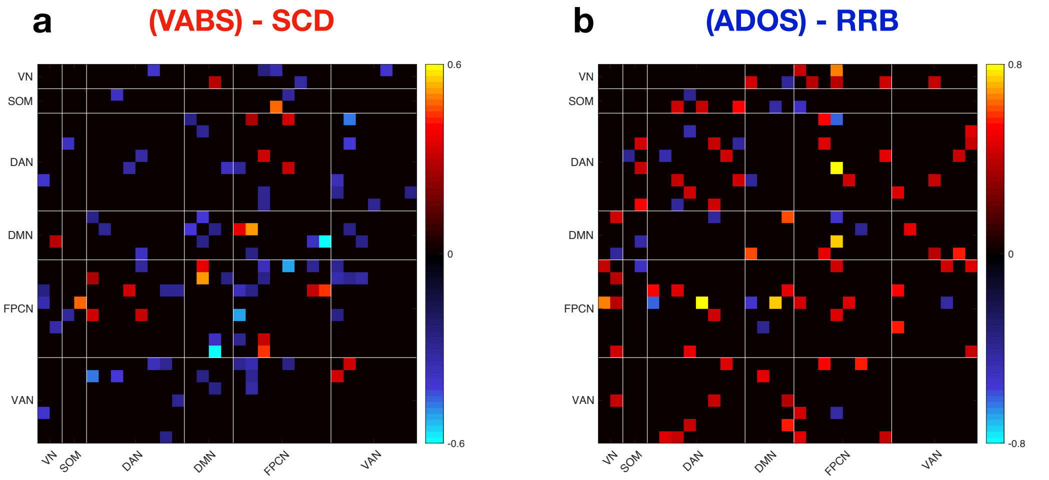


**Supplementary Figure 7.** Composition of ROI-level FC for symptom-linked FC dimensions. **a** SCD-linked dimension. The hypo-connectivity between left angular gyrus (in DMN) and right middle temporal gyrus (in FPCN) shows strongest contribution to SCD-linked FC dimension score. **b** RRB-linked dimension. The hyper-connectivity between left frontal eye field (in DAN) and right inferior parietal lobe (in FPCN) shows strongest contribution to RRB-linked FC dimension score.

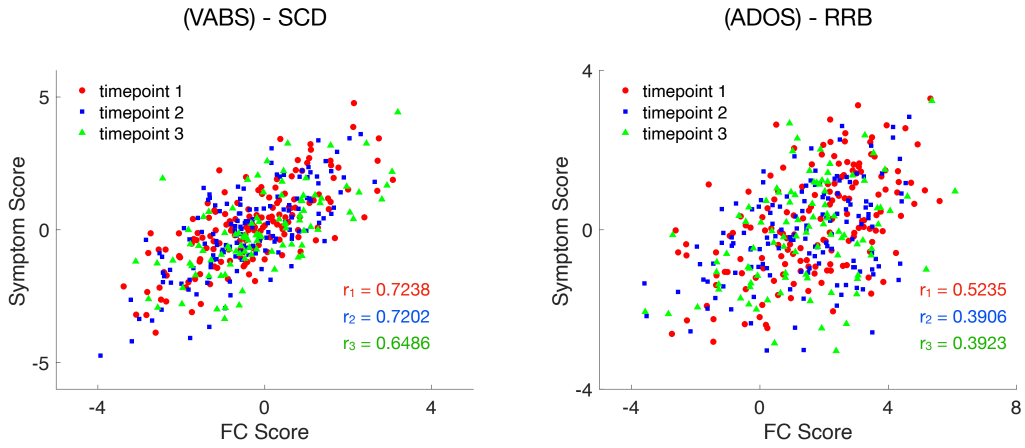


**Supplementary Figure 8.** Associations between symptom and FC dimension scores are general to longitudinal assessments. Fisher's z test for highest and lowest correlations: SCD: $p = 0.28$; RRB: $p = 0.15$. p-values are uncorrected.

**Supplementary Table 1** Distribution of Demographics and Psychiatric Assessments of ASD Patients in ABC-CT Dataset

| N = 392 | | Metrics | | | |
|---|---|---|---|---|---|
| Male: 289 (73.7%) Female: 103 (26.3%) | | Mean | Median | Min | Max |
| Demographics | Age | 8.83 | 8.71 | 6.08 | 12.08 |
| ADOS | SA | 9.47 | 9 | 3 | 19 |
| | RRB | 3.55 | 3 | 0 | 8 |
| AIM | Freq_RB | 19.37 | 18 | 8 | 40 |
| | Freq_Comm | 9.49 | 9 | 5 | 24 |
| | Freq_SRc | 18.04 | 18 | 7 | 34 |
| | Freq_AtB | 11.61 | 11 | 5 | 25 |
| | Impact_RB | 15.77 | 14 | 8 | 40 |
| | Impact_Comm | 8.73 | 8 | 5 | 25 |
| | Impact_SRc | 12.80 | 11 | 7 | 35 |
| | Impact_AtB | 9.79 | 9 | 5 | 25 |
| SRS | SAwr | 69.43 | 70 | 15 | 100 |
| | SCog | 68.32 | 68.5 | 16 | 104 |
| | SComm | 69.87 | 70.5 | 26 | 106 |
| | SMotiv | 62.39 | 62 | 14 | 106 |
| | RRB | 70.62 | 71 | 17 | 108 |
| VABS | Comm_1 | 10.53 | 10 | 1 | 18 |
| | Comm_2 | 11.26 | 11 | 1 | 18 |
| | Comm_3 | 13.16 | 13 | 5 | 24 |
| | Social_1 | 10.94 | 11 | 1 | 19 |
| | Social_2 | 11.20 | 11 | 2 | 21 |
| | Social_3 | 12.73 | 13 | 4 | 21 |
| | DL_1 | 9.24 | 10 | 1 | 18 |
| | DL_2 | 11.04 | 12 | 1 | 22 |
| | DL_3 | 10.58 | 10 | 3 | 21 |

**Supplementary Table 2** Distribution of Demographics and Psychiatric Assessments of ASD Patients in HBN Dataset

| N = 222 | | Metrics | | | |
|---|---|---|---|---|---|
| Male: 185 (83.3%) | Female: 37 (16.7%) | Mean | Median | Min | Max |
| Demographics | Age | 10.97 | 10.21 | 5.85 | 21.48 |
| SRS | SAwr | 65.98 | 67 | 34 | 90 |
| | SCog | 68.96 | 68 | 40 | 90 |
| | SComm | 69.62 | 70 | 41 | 90 |
| | SMotiv | 64.60 | 62 | 40 | 90 |
| | RRB | 70.48 | 71 | 42 | 90 |
| RBS | SB | 3.76 | 2 | 0 | 21 |
| | SIB | 1.29 | 1 | 0 | 14 |
| | CB | 1.95 | 1 | 0 | 12 |
| | RSB | 7.53 | 5 | 0 | 34 |
| | RI | 2.42 | 2 | 0 | 9 |

Abbreviations: **SB**: Stereotypical Behavior; **SIB**: Self-Injurious Behavior; **CB**: Compulsive Behavior; **RSB**: Ritualistic/Sameness Behavior; **RI**: Restricted Interests.
